# SSR1 is a vital regulator in plant mitochondrial iron-sulfur biosynthesis

**DOI:** 10.1101/2021.03.09.434627

**Authors:** Xuanjun Feng, Huiling Han, Diana Bonea, Jie Liu, Wenhan Ying, Yuanyuan Cai, Min Zhang, Yanli Lu, Rongmin Zhao, Xuejun Hua

## Abstract

The Arabidopsis *SHORT AND SWOLLEN ROOT1* (*SSR1*) gene encodes a mitochondrial TPR domain-containing protein and was previously reported to function in maintaining mitochondria function. In a screen for suppressors of the short-root phenotype of the loss-of-function mutant *ssr1-2*, two mutations, *sus1* and *sus2* (suppressor of *ssr1-2*), were isolated. *sus1* and *sus2* result from G87D and T55M single amino acid substitution in HSCA2 (At5g09590) and ISU1 (At4g22220), both of which are core components in iron-sulfur cluster biosynthesis pathway in mitochondria (ISC). We here demonstrated that SSR1 displayed a strong chaperone-like activity and was able to enhance the binding of HSCA2 to ISU1, an essential step for the normal operation of ISC machinery. Accordingly, the enzymatic activities of several iron-sulfur proteins, the mitochondrial membrane potential and ATP content are reduced in *ssr1-2*. Interestingly, *SSR1* appears to exist only in plant lineages, possibly conferring adaptive advantages on plant ISC machinery to environment.

## Introduction

Iron-sulfur (Fe-S) cluster is a cofactor for many enzymes that play vital roles in biological processes like respiration, photosynthesis, DNA repair and hormone synthesis (Balk and Pilon, 2011; Balk and Schaedler, 2014). The cellular biosynthesis and assembly of Fe-S cluster is evolutionarily conserved from bacteria to higher plants and mammals (Beinert, 2000; Barras et al., 2005). Three Fe-S cluster biosynthesis pathways have been reported in bacteria, namely ISC, SUF and NIF (Johnson et al., 2005; Ayala-Castro et al., 2008; Fontecave and Ollagnier-de-Choudens, 2008; Roche et al., 2013). NIF pathway exists only in nitrogen-fixing bacteria *A. vinelandii* while SUF and ISC pathways are conserved in most prokaryotes (Roche et al., 2013). In higher organisms, the two organelles, plastids and mitochondria, have inherited the SUF and ISC pathways from their endosymbiotic ancestors, cyanobacteria and proteobacteria, respectively.

The ISC pathway has been well-characterized in bacteria and yeast. In bacteria, five core proteins have been identified: IscS, IscU, Fdx, HscB and HscA (Roche et al., 2013). The yeast homologs are Nfs1, Isu1, Yfh1, Jac1, and Ssq1 (Barras et al., 2005; Dutkiewicz et al., 2017). During the Fe-S assembling stage, IscS/Nfs1 catalyzes the release of sulfur from L-cysteine and transfers it to the scaffold protein IscU/Isu1, where Fe-S clusters are assembled (Smith et al., 2001; Urbina et al., 2001; Cupp-Vickery et al., 2003; Smith et al., 2005). Fdx/Yfh1 was proposed to supply iron to scaffold IscU/Isu1 (Chandramouli et al., 2007). Once assembled, Fe-S clusters are released from IscU/Isu1 and transferred to recipient apoproteins. During this stage, HscA/Ssq1 and HscB/Jac1, members of the DnaK family and J-domain protein family chaperones, respectively, are required (Hoff et al., 2000). HscA/Ssq1 recognizes the LPPVK motif in IscU/Isu1 to form a protein complex together with apoproteins (Hoff et al., 2003; Cupp-Vickery et al., 2004). IscU/Isu undergoes a conformation change to facilitate the release of Fe-S cluster from IscU/Isu1 (Kim et al., 2012; Balk and Schaedler, 2014; Dutkiewicz et al., 2017). This process is energized by HscA ATP hydrolysis, which is generally stimulated by the binding of HscB/Jac1 and/or IscU/Isu1 (Dutkiewicz et al., 2004; Kim et al., 2012; Majewska et al., 2013; Leaden et al., 2014). In addition, the binding of HscB/Jac1 to IscU/Isu1 promotes the disassociation of IscS/Nsf1 from IscU/Isu1, and the association of HscA/Ssq1 to IscU/Isu1 (Hoff et al., 2003; Cupp-Vickery et al., 2004; Kim et al., 2012; Kim et al., 2012; Majewska et al., 2013).

Many Arabidopsis genes encoding the core components of ISC pathway have been identified based on sequence homology with their counterparts in bacteria or yeast, and some core proteins are encoded by a multiple-gene family (Tone et al., 2004; Leon et al., 2005; Frazzon et al., 2007; Xu et al., 2009; Balk and Schaedler, 2014). For instance, there are three genes for cysteine desulfurase in Arabidopsis, namely *NFS1, NFS2* and *ABA3* (Balk and Schaedler, 2014; Armas et al., 2020). Only NFS1 showed mitochondria localization and high homology with IscS, a bacterial cysteine desulfurase (Frazzon et al., 2007). Scaffold protein ISU is also encoded by three genes, *ISU1, ISU2* and *ISU3*, in Arabidopsis, with *ISU1* being expressed most abundantly (Leon et al., 2005; Frazzon et al., 2007). However, the mechanistic functions of the individual isoform still remain elusive. Moreover, like those in bacteria, Arabidopsis HSCA1, HSCA2, HSCB and ISU1 could interact with each other, and the binding with HSCB and/or AtISU1 also promotes the ATPase activity of HSCA2 (Leaden et al., 2014). Additionally, Arabidopsis HSCB was found to be able to rescue the phenotype of the yeast *Jac1* mutant, though it is localized in both mitochondria and cytosol (Xu et al., 2009).

Despite the identification and extensive study of plant ISC pathway genes, whether plants possess any specific regulatory mechanisms is still unknown. Previously, we have characterized an Arabidopsis gene *SHORT AND SWOLLEN ROOT1* (*SSR1*), encoding a mitochondrial protein with TPR domain, for its roles in regulating root growth. Loss-of-function *ssr1* mutants display dramatically shortened roots (Zhang et al., 2015) and impaired mitochondrial function (Han et al., 2021). Here, by screening and analyzing the suppressors of *ssr1-2*, we provided evidence that SSR1, acting as a chaperone-like component in the Arabidopsis ISC pathway, interacts with both ISU1 and HSCA2 to strengthen their interaction. This would likely facilitate the release of Fe-S cluster from ISU1 scaffold. In addition, *SSR1* is uniquely present in plant species and its homolog could not be found in organisms of other kingdoms. We, therefore, propose that *SSR1* is a plant-specific molecular chaperone to critically regulate the ISC pathway in mitochondrial Fe-S cluster biosynthesis.

## Results

### *sus1* and *sus2* suppress the short-root phenotype of *ssr1-2*

By analyzing a T-DNA insertional mutant *ssr1-2*, we have previously shown that knockout of the *SSR1* gene resulted in a severe root growth defect phenotype (Zhang et al., 2015). To better understand the molecular mechanism of *SSR1* in controlling root development, the *ssr1-2* mutant was mutagenized by ethyl methanesulfonate and the M2 progenies were used for screening the suppressors of the short-root phenotype of *ssr1-2*. From a dozen identified suppressors, two lines, designated as *sus1* and *sus2* (*<u>su</u>ppressor* of *<u>s</u>sr1-2*), have the best recovered root length (about 85% and 100%, respectively) at seedling stage as well as a complete rescue of the dwarf phenotype at the flowering stage (Fig. 1A).

**Figure 1.**
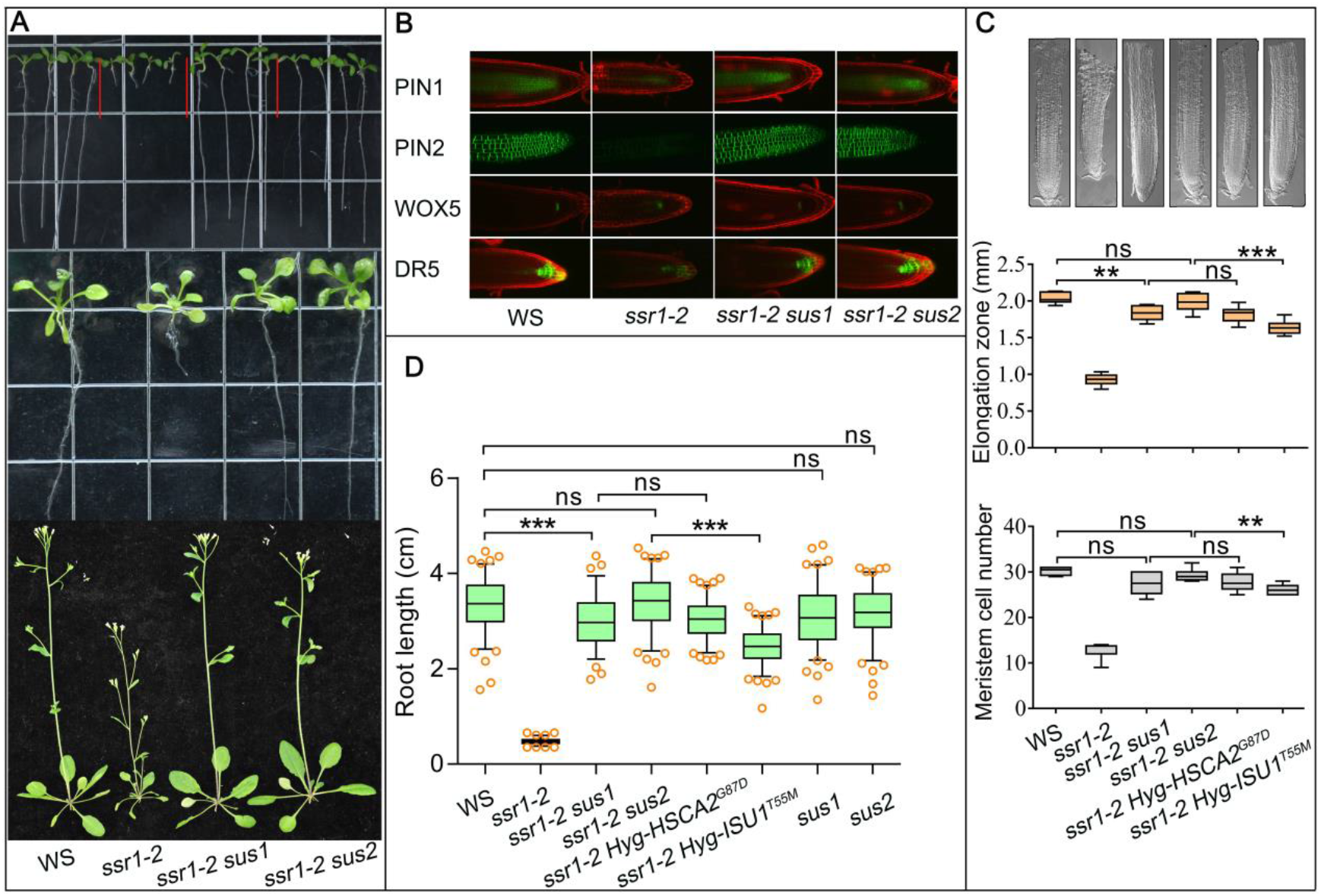
Identification of *ssr1-2* suppressor mutant genes *sus1* (*HSCA2*^*G87D*^) and *sus2* (*ISU1*^*T55M*^). Two suppressors (*sus1* and *sus2*) were identified to rescue the growth defect of *ssr1-2* and the whole genome sequencing identified *sus1* as *HSCA2*^*G87D*^ while *sus2* as *ISU1*^*T55M*^. (*A*) *sus1* and *sus2* rescued both the root and shoot growth defects of *ssr1-2* to the levels comparable to wild type (WS). Top, 10-day-old seedlings; middle, 20-day-old seedlings; bottom, 40 -day-old plants. (*B*) *sus1* and *sus2* rescued the expression defects in *ssr1-2* of auxin transport and response marker genes *PIN1::PIN1GFP, PIN2::PIN2GFP, WOX5::WOX5GFP* and *DR5::GFP*, labelled as PIN1, PIN2, WOX5, and DR5 respectively, to the levels comparable to those in WS. (*C*) *sus1* and *sus2* mutant genes that were cloned, designated as *HSCA2*^*G87D*^ and *ISU1*^*T55M*^, respectively, and re-transformed back to *ssr1-2* plants, rescued the swollen short root phenotype. Top, images of primary roots of different lines at 10-day-old. Middle, elongation zone length, bottom, meristem zone cell number. In transgenic lines, the transgenes are associated with a hygromycin resistant gene and therefore designated as *Hyg-HSCA2*^*G87D*^ or *Hyg-ISU1*^*T55M*^. It should be noted that *ssr1-2 Hyg-HSCA2*^*G87D*^ still contains wild type *SUS1* allele and *ssr1-2 Hyg-ISU*^*T55M*^ contains wild type *SUS2* allele. (*D*) primary root length of different lines grown for 10 days. Lines analyzed are the same as in (*C*) except that *HygR-HSCA2*^*G87D*^ and *HygR-ISU1*^*T55M*^ represent the lines that have cloned *sus1* gene *HSCA2*^*G87D*^ and *sus2* gene *ISU1*^*T55M*^ re-transformed into WS wild type background, respectively. In (*C*) and (*D*), 15 and 100 roots of each were used for statistical analysis by ANOVA (in *C*, for elongation zone, *F*=123.2, *DFn*=5, for stem cell number, *F*=118, *DFn*=5; in d, *F*=542.8, *DFn*=7). statistical value is shown as box & whiskers with 5-95 percentile. Asterisk “*”, “**” and “***” represent statistics analysis with *p* values < 0.05, 0.01, 0.001 respectively. “ns” represents statistically not significant.

It has been previously reported that several auxin-related markers were abnormally expressed in *ssr1-2* seedlings (Zhang et al., 2015). To examine whether *sus1* and *sus2* also rescued the abnormal expression of the auxin-related genes, *PIN1-GFP, PIN2-GFP, Wox5-GFP* and *DR5-GFP* marker lines in *ssr1-2* background were crossed to double mutants *ssr1-2 sus1* and *ssr1-2 sus2*. The resulting F3 progenies which are homozygous for both *sus* and marker genes were identified. Confocal microscopic analyses showed that the expression patterns of all four auxin transport or signaling related genes in double mutants were restored and resembling those in the wild type background (Fig. 1B). These data indicate that *sus1* and *sus2* are two suppressor mutants that could indeed rescue the defects in root growth and structure of *ssr1-2* and therefore were chosen for further analysis.

To investigate the genetic nature of *sus1* and *sus2*, we backcrossed the two suppressor mutants (designated as *ssr1-2 sus1* and *ssr1-2 sus2* respectively) with *ssr1-2*. All *ssr1-2 sus1/SUS1* F1 seedlings crossed from *ssr1-2 sus1* parent exhibited similar root length to *ssr1-2 sus1* suggesting that *sus1* is a dominant mutation. In contrast, the primary roots of *ssr1-2 sus2/SUS2* F1 seedlings crossed from *ssr1-2 sus2* parent are longer than that of *ssr1-2*, but less pronounced compared to *ssr1-2 sus2*, suggesting that *sus2* is a semi-dominant mutation. Further analysis of self-pollinated F2 progenies confirmed these observations, and a clear 3:1 ratio and 1:2:1 ratio were observed for *sus1* and *sus2* mutations, respectively, in term of their root lengths (Supplemental Fig. S1 and Table S1).

### *SUS1* and *SUS2* encode mitochondrial chaperone protein HSCA2 and Fe-S cluster assembly protein ISU1

After twice backcrosses to parent *ssr1-2* and super bulked segregant analysis, five candidate genes from each suppressor line were identified, and then verified by genetic complementation. Since *sus1* and *sus2* are dominant and semi-dominant mutations, respectively, the complementation analysis was performed by re-introducing the full-length genomic fragments of candidate genes from the suppressor lines into *ssr1-2*. It turned out that re-introducing *At5g09590* from *ssr1-2 sus1* and *At4g22220* from *ssr1-2 sus2* could rescue the root apical meristem cell division and elongation zone growth defect (Fig. 1C), as well as the primary root length phenotype of *ssr1-2* (Fig. 1D), thus confirming that *At5g09590* and *At4g22220* are *SUS1* and *SUS2*, respectively. *At5g09590* and *At4g22220* encode a mitochondrial heat shock cognate 70 (mtHSC70-2 or HSCA2) and an ISU1 protein, respectively, both being previously reported and critically involved in Fe-S cluster assembly in mitochondria (Balk and Schaedler, 2014; Leaden et al., 2014). *sus1* and *sus2* each bears a point substitution mutation in the coding region of corresponding *HSCA2* and *ISU1* genes. *sus1* carries a G to A transition at position 260, resulting in an HSCA2^G87D^ mutant protein. *sus2* carries a C to T transition at position 164, resulting in an ISU1^T55M^ mutant protein.

With the identification of the two suppressor genes which cause amino acid substitution in HSCA2 (HSCA2^G87D^) and ISU1 (ISU1^T5M^), we took a different approach to analyze the other candidate suppressors obtained from our initial screen. The genomic sequences of *HSCA2* and *ISU1* genes from *sus4* to *sus8* suppressor lines were all cloned and sequenced. Interestingly, *sus4, sus5, sus6, sus7* and *sus8* mutants all carry point mutation in *HSCA2* or *ISU1*, resulting in mutant protein ISU1^A143V^, ISU1^G106D^, ISU1^A143T^, HSCA2^R394C^ and ISU1^A140V^, respectively. Their suppressor function was partly verified by genetic complementation with two representative suppressor genes *sus5* (encoding ISU1^G106D^) and *sus6* (encoding ISU1^A143T^), both indeed rescuing the short-root phenotype of *ssr1-2* (Fig. 2A).

**Figure 2.**
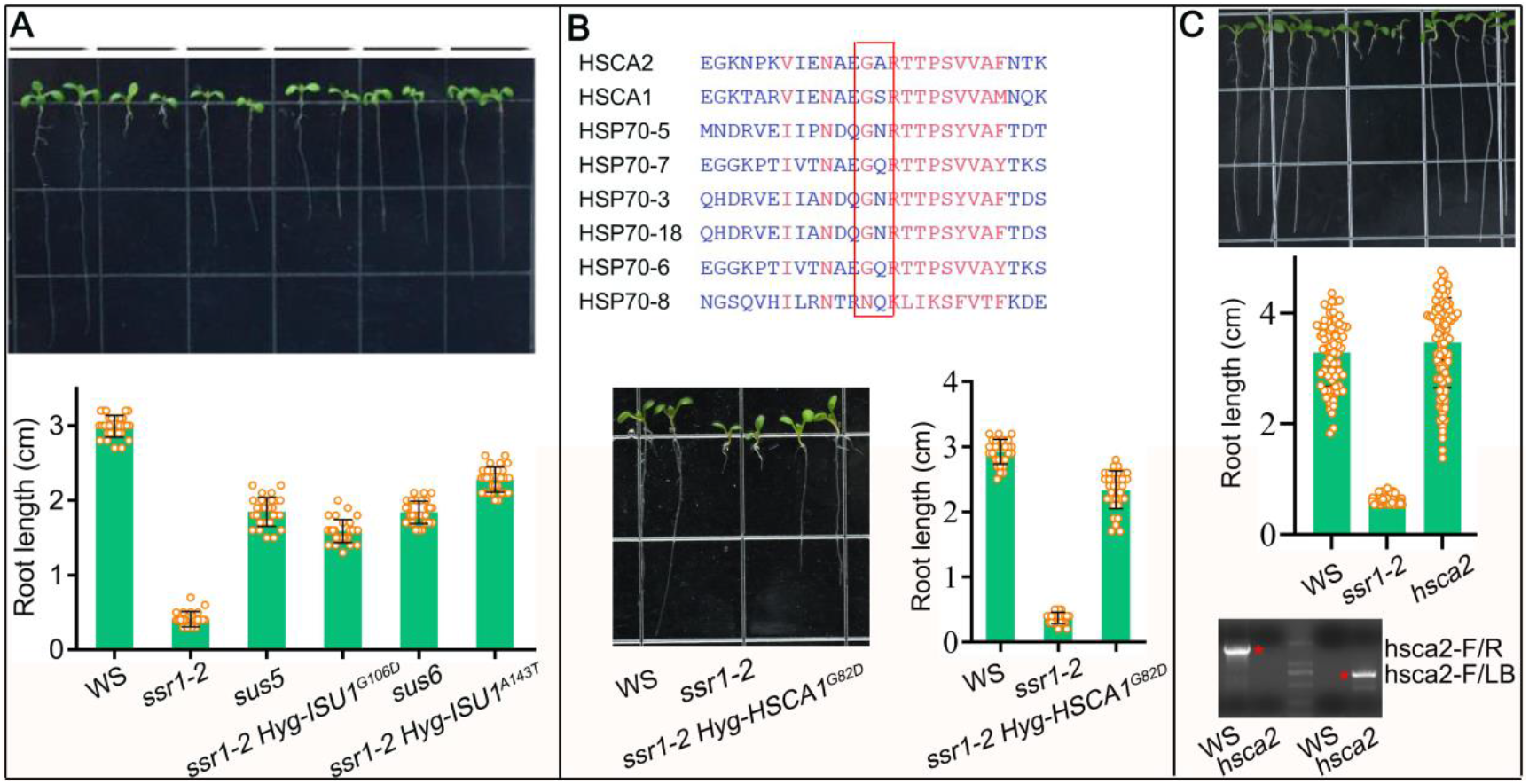
Effects of ISU1^G106D^, ISU1^A143T^, HSCA1^G82D^ point mutations and *hsca2* knockout mutant on root growth. (*A*) *ISU1G*^*106D*^ and *ISU1*^*A143T*^ are *SUS5* and *SUS6* respectively, and they can partially rescue the phenotypes of *ssr1-2*. (*B*) HSCA1^G82D^ is a homolog of HSCA2^G87D^, and it can rescue the phenotypes of *ssr1-2*. (*C*) HSCA2 loss of function mutation has no effect on root length. Primers hsca2-F and hsca2-R were used for the amplification of *HSCA2* genome fragment, and hsca2-F and LB were used to detect the T-DNA insertion. For (*A-C*), representative seedlings grown for 10 days and average root length of analyzed lines were shown. Error bars represent standard deviation from 20 (*A*), 30 (*B*), and 60 (*C*) seedlings on primary roots.

HSCA1 is a homolog of HSCA2, and also located in mitochondria (Balk and Schaedler, 2014). To test whether HCSA1 has an overlapping function with HSCA2, we cloned and constructed an HSCA1^G82D^ mutant which corresponds to HSCA2^G87D^ and introduced it into *ssr1-2* under the native promoter. The primary root length of *ssr1-2* seedlings was substantially increased with the expression of HSCA1^G82D^ (Fig. 2B), suggesting that HSCA1 function may partially overlap with that of HSCA2 in vivo. Furthermore, we obtained and confirmed a knock-out mutant of *HSCA2, hsca2* (CS479451 from ABRC), and interestingly *hsca2* seedlings did not show any significant phenotype regarding primary root length (Fig. 2C). All these data clearly indicate HSCA1 and HSCA2 are functionally redundant, at least in controlling root development under normal growth conditions. This also agrees with the common view that, after the duplication event of certain mitochondrial HSP70 gene, one copy, HSCA1 in Arabidopsis, is the predominant form and plays a multifaceted role, while the paralog HSCA2 becomes an isoform only for the ISC pathway (Dutkiewicz et al., 2017).

### SSR1, ISU1 and HSCA2 function collaboratively in regulating root growth

Next, we determined to explore the molecular mechanism underlying suppression of shoot-root phenotype of *ssr1-2*. First, we would like to understand whether *sus1* or *sus2* could affect root growth in WT background. For this analysis, *sus1* and *sus2* single mutants were isolated by back-crossing *ssr1-2 sus1* and *ssr1-2 sus2* with the wild-type WS and their root length was characterized. Under the normal growth conditions, the root length of *sus1* and *sus2* single mutants was similar to that of the wild type (Fig. 1D), implying that the identified suppressors are specifically functioning in the root growth process affected by *SSR1* mutation.

Subsequently, we set to investigate the tissue-specific expression of *SSR1, HSCA2* and *ISU1* to see if their expression patterns overlap with one another. It has been previously reported that *ISU1* is highly expressed in root and leaf throughout of the life cycle (Leon et al., 2005; Frazzon et al., 2007; Tsugama et al., 2009). We then analyzed the expression patterns of *HSCA2* and *SSR1* by fusing their promoters to a *GUS* reporter gene. *SSR1* promoter activity was generally weak in most tissues as analyzed at the seedling and flowering stages but could be well detected in root tips (Fig. 3A). *HSCA2* promoter displayed high activity in aerial parts and an unexpectedly low activity, but still detectable, in root tips (Fig. 3B).

**Figure 3.**
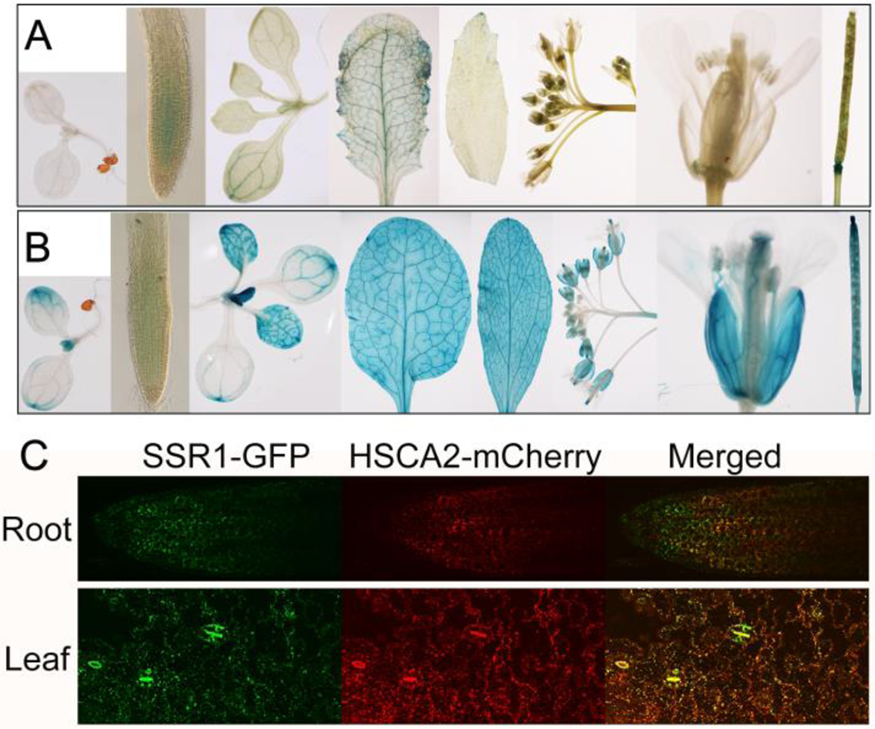
Tissue specific expression of *SSR1* and *HSCA2*. The expression of GUS driven by promoters of *SSR1* (*A*) and *HSCA2* (*B*) in various organs. (*C*) The expression of *SSR1-GFP* and *HSCA2-mCherry* driven by native promoters in root tips and cotyledons.

To better understand how HSCA2 and SSR1 are co-expressed in the mitochondria, HSCA2 and SSR1 were fused with *mCherry* and *GFP* reporter genes, respectively, and introduced into Arabidopsis plants under their native promoters. HSCA2-mCherry and SSR1-GFP expressions were well observed in root tips and cotyledons, and co-localized as discrete spots representing mitochondria (Fig. 3C, visible as yellow dots). Therefore, our results and the results of others (Frazzon et al., 2007) indicated that SSR1, ISU1 and HSCA2 are indeed co-expressed in mitochondria, hence, supported that they may act collaboratively in affecting root growth.

We conducted a comprehensive database search for all *SSR1* homologs, and the bioinformatics analysis indicated that SSR1 homolog is present in green algae, moss, lower plants and the higher plant species most as a single copy, but absent in bacteria, fungi and animalia (Supplemental Fig. S2). Although it is unknown whether SSR1 in other species plays a similar role, it is obvious that it is required to regulate certain metabolic pathways only in plant mitochondria.

### SSR1 interacts with HSCA2 and ISU1 and promotes HSCA2-ISU1 association

To further understand how *SSR1* is involved in the ISC pathway, we analyzed possible protein-protein interactions between SSR1, HSCA2 and ISU1 by biomolecular fluorescence complementation (BiFC) assay using Arabidopsis mesophyll protoplasts. Fluorescence signals were well visualized in protoplasts as discrete bright spots, reminiscent to cellular mitochondria, when cCFP-tagged SSR1 was co-expressed with nVenus-tagged HSCA2 or ISU1 (Fig. 4A). This indicates that both HSCA2 and ISU1 physically interact with SSR1. However, the interaction between ISU1 and SSR1 seems weaker than that between HSCA2 and SSR1 (Fig. 4A).

**Figure 4.**
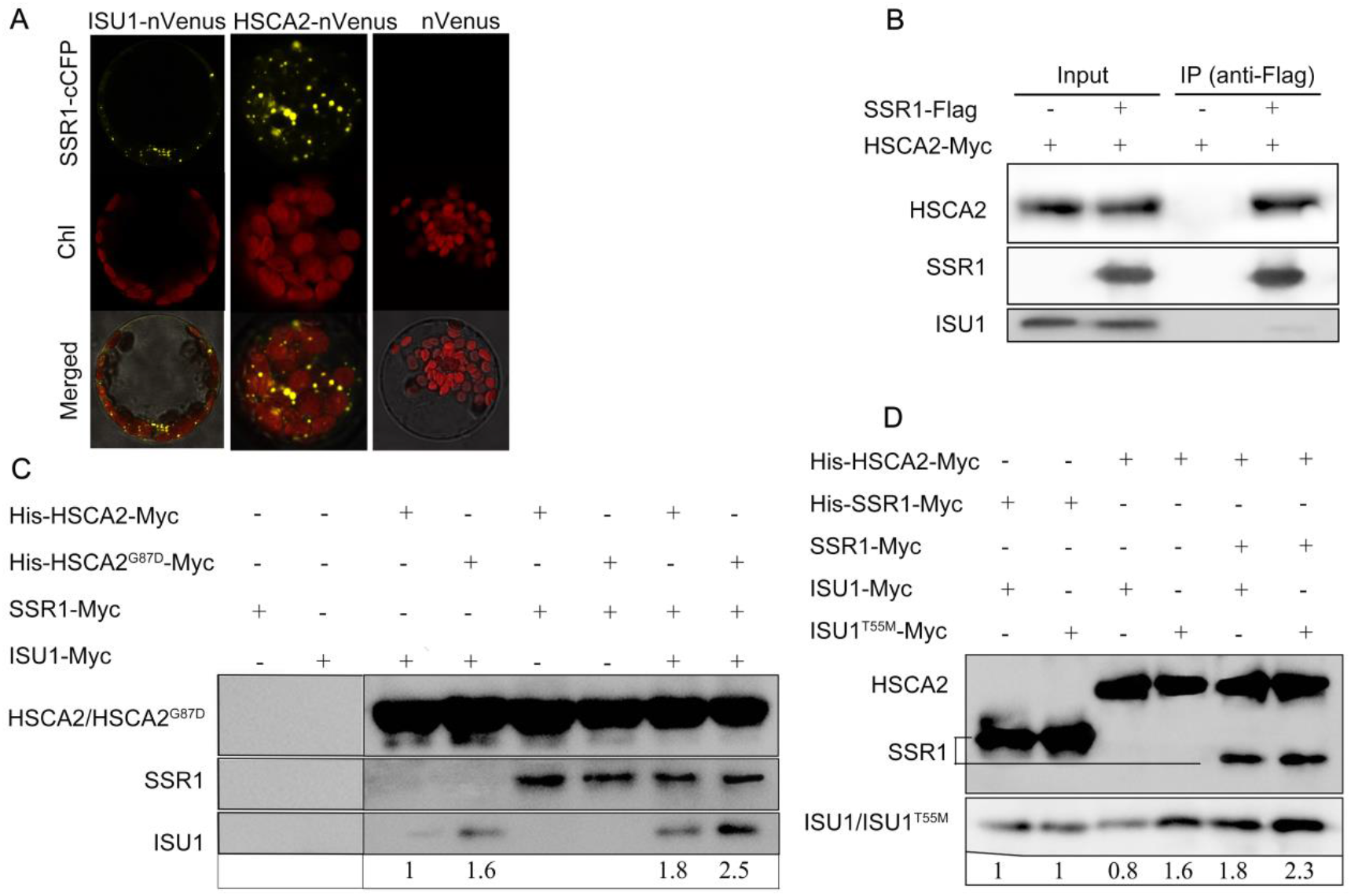
SSR1 interacts with HSCA2 and ISU1 and promotes HSCA2-ISU1 association. (*A, B*) *in vivo* interactions between SSR1 and HSCA2 or ISU1 were analyzed in Arabidopsis protoplasts by BiFC (*A*) and co-immunoprecipitation (*B*) assays. In (*A*), SSR1 was fused to C-terminal half of CFP (SSR1-cCFP) and HSCA2 or ISU1 was fused to the n-terminus of Venus (HSCA2-nVenus, ISU1-nVenus). Chlorophyll autofluorescence (chl) was also shown. In (*B*), stable transgenic lines co-expressing Flag-tagged SSR1 under CaMV 35S promoter and cMyc-tagged HSCA2 under its native promoter were immunoprecipitated with anti-Flag antibody and co-purified proteins were immunodetected with anti-Myc or anti-ISU antibodies. (*C, D*) in vitro pull-down assays using Ni-NTA resin with His6-tagged proteins expressed in *E. coli* as baits. Co-purified proteins were detected with anti-Myc antibody. The immunoblotting signals of ISU1-Myc or ISU1^T55M^-Myc in (*C*) and (*D*) were quantified by ImageJ and relative intensities were shown under respective lanes.

To further confirm the interaction, co-IP assays were performed using transgenic plants. Since the activity of native *SSR1* promoter was rather weak, *SSR1-Flag* construct driven by 35S promoter was used and co-integrated into *ssr1-2* mutants with *HSCA2-Myc* driven by native promoter via transformation and subsequent hybridization. Protein extracts from transgenic plants were incubated with anti-Flag antibody resin to purify protein complexes containing SSR1-Flag. Subsequent immunoblotting analysis with anti-Myc or anti-ISU1 antibodies demonstrated that both HSCA2-Myc and ISU1 were co-purified with SSR1-Flag (Fig. 4B), though with much less ISU1 being co-purified compared with HSCA2. This was consistent with the BiFC assay (Fig. 4A) and indicated that SSR1 interacts much weaker with ISU1 than with HSCA2.

Since ISU1 is known to interact with HSCA2 *in vivo* (Balk and Schaedler, 2014; Leaden et al., 2014), the above co-IP assay with SSR1-Flag as a bait cannot rule out the possibility that SSR1 interacts with ISU1 or HSCA2 indirectly. Therefore, interactions of these three proteins and two mutant forms, HSCA2^G87D^ and ISU1^T55M^, were further investigated by in vitro pull-down assays using proteins expressed and purified from *E. coli*. The protein complexes containing His_6_-HSCA2-Myc or His_6_-HSCA2^G87D^-Myc were pulled down by Ni-Sepharose and analyzed by Western blot with anti-Myc antibody. It is evident that SSR1-Myc can be co-purified with His_6_-HSCA2-Myc or His-HSCA2^G87D^-Myc (Fig. 4C), indicating that SSR1 and HSCA2 directly interact with each other.

To confirm the physical interaction between SSR1 and ISU1, another pull-down assay using His_6_-SSR1-Myc as a bait was performed. ISU1-Myc and ISU1^T55M^-Myc can be co-purified with SSR1, suggesting SSR1 also interacts directly with ISU1 (Fig. 4D). Interestingly, the association between ISU1 and HSCA2^G87D^ was significantly stronger than that between ISU1 and HSCA2 (Fig. 4C). Similarly, HSCA2 displayed stronger interaction with ISU1^T55M^ than with ISU1 (Fig. 4D). Additionally, the presence of SSR1 further promoted the interaction between ISU/ISU1^T55M^ and HSCA2/HSCA2^G87D^ (Fig. 4, C and D).

Based on these ternary protein-protein interactions (Fig. 4), we speculated that the enhanced affinity between ISU1 and HSCA2, when a point mutation in either ISU1 (ISU1^T55M^) or HSCA2 (HSCA2^G87D^) is present, may be responsible for the suppression of the root growth defect in *ssr1-2*. To verify this hypothesis, amino acid substitutions were introduced into ISU1^T55M^ at the L^126^PPVK^130^ motif, which was previously reported to mediate the interaction with HSCA (Hoff et al., 2003; Cupp-Vickery et al., 2004; Dutkiewicz et al., 2004). In vitro pull-down assays showed that L126A, P127A, or P128S mutation slightly impaired the interaction between ISU1^T55M^ and HSCA2, while V129E or K130A mutation dramatically reduced the affinity between ISU1^T55M^ and HSCA2 (Fig. 5A). In addition, simultaneous substitution of PVK to AAA almost completely abolished the interaction between ISU1^T55M^ and HSCA2 (Fig. 5A). Subsequently, the ISU1 mutant constructs were introduced into *ssr1-2* plant to test their ability to suppress the short-root phenotype of *ssr1-2*. It is interesting to note that ISU1^T55ML126A^, ISU1^T55MP127A^, and ISU1^T55MP128S^, but not ISU1^T55MV129E^, ISU1^T55MK130A^, and ISU1^T55MAAA^ partially rescue *ssr1-2* (Fig. 5B), well in agreement with their ability to interact with HSCA2. Taken all these together, we have demonstrated that enhanced interaction between ISU1 and HSCA2 is essential for the rescue of *ssr1-2* phenotype.

**Figure 5.**
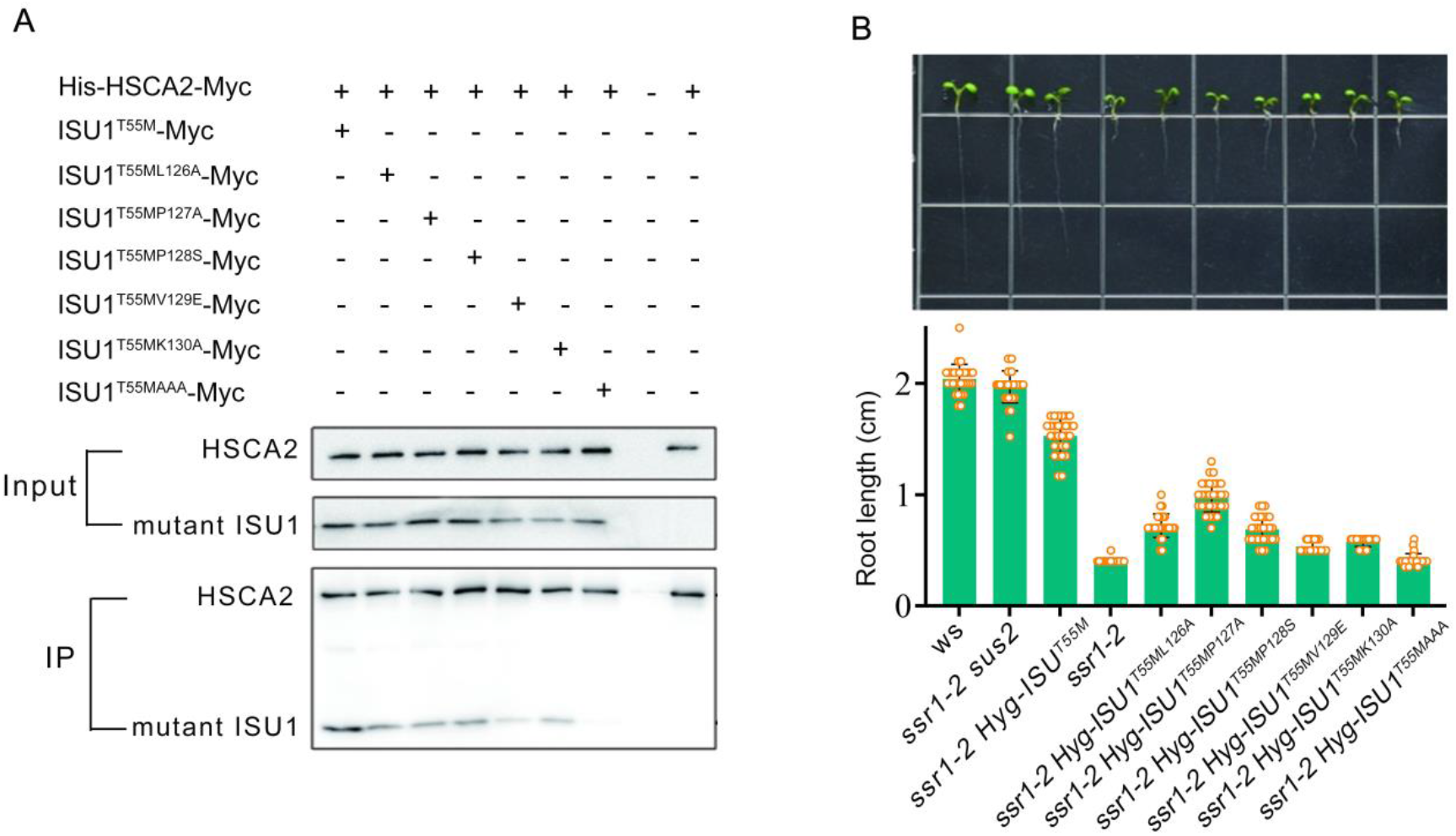
Enhanced interaction between ISU1 and HSCA2 is essential for the rescue of *ssr1-2* growth defect. (*A*) Ni-NTA in vitro pull-down assays with His6-tagged HSCA2 expressing in *E. coli* as the bait. Different Myc-tagged ISU1 mutation variants within the LPPVK motif were expressed in *E. coli* and differentially co-purified with HSCA2. (*B*) root length of wild type (WS) and *ssr1-2* mutants carrying *sus2* allele or transformed with an additional ISU1 gene but with the LPPVK motif differentially mutated. Top, representative seedlings at 8-days-old. Bottom, root lengths shown as bar graphs with error bars representing standard deviations from 20 seedlings.

### SSR1 displays a chaperone-like function

The involvement of HscA/HscB chaperone system in the assembly of Fe-S clusters has been well documented in bacteria, yeast and plant (Hoff et al., 2000; Roche et al., 2013; Balk and Schaedler, 2014; Leaden et al., 2014). The release of mature Fe-S cluster from ISU1 requires the ATP binding and hydrolysis activity of chaperone HscA, and this process can be stimulated by HscB (Hoff et al., 2000). Since we have shown that SSR1 interacts with HSCA2 and ISU1 both *in vivo* and *in vitro*, we speculated that SSR1 may display HscB/Jac1-like activity. To explore this possibility, we used purified SSR1, HSCA2 and HSCA2^G87D^ from *E. coli* to test their general chaperone activity in preventing heat-induced substrate protein from aggregation.

By using citrate synthase (CS) as a model substrate, it was shown that both HSCA2 and HSCA2^G87D^ have a strong general chaperone activity, and the difference between the wild-type and the mutant form is subtle (Fig. 6A and Supplemental Fig. S3). ISU1 itself surprisingly inhibited heat-induced aggregation of CS though not as efficiently as HSCA2 (Fig. 6B). When ISU1 and HSCA2/HSCA2^G87D^ were both added with CS, much more aggregates accumulated especially at the later time (Fig. 6, B and C). This implied that certain physical interactions occurred between HSCA2 and ISU1. Interestingly, whenever SSR1 is present, no aggregate formed, indicating SSR1 has a powerful chaperone-like activity (Fig. 6, B and C). Further titration analysis showed that SSR1 itself displayed a very strong chaperone activity in preventing heat-induced citrate synthase (CS) from aggregating even at a very low concentration (Fig. 6D). Although the general chaperone activity assay may not be sufficient in revealing detailed structural features of the tested proteins, it is plausible to propose that HSCA2, ISU1, and SSR1 can form a protein complex in which one or more proteins have undergone a significant conformational change.

**Figure 6.**
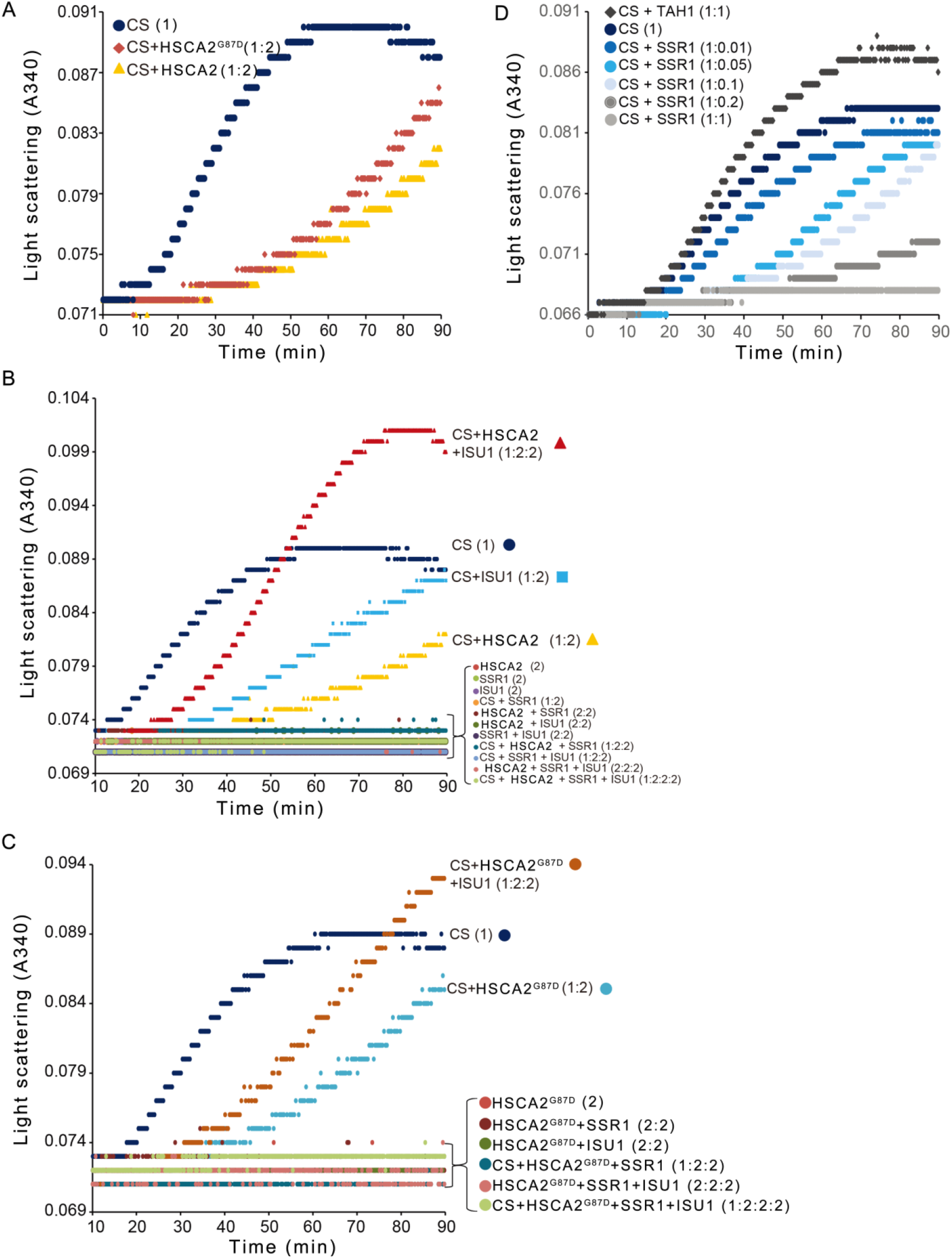
SSR1 appears strong general molecular chaperone activity. Heat-induced aggregation of citrate synthase (CS) was performed at 45°C for 90min and monitored by increased light scattering at 340nm. The molecular ratios of CS to tested proteins are indicated within brackets following each protein sample. Control samples without adding CS or those tested protein mixtures that did not show significant protein aggregation are showing curves crowded along the basal line, and therefore shown together without referring individual curve. Only one representative absorbance curve is shown for each tested sample mixture. (*A*) Heat-induced CS aggregation with wildtype HSCA2 (wtHSCA2) and mutant form HSCA2 (mtHSCA2). (*B*) Heat-induced CS aggregation with wildtype HSCA2 (wtHSCA2) in combination with ISCU1 and/or SSR1. (*C*) Heat-induced CS aggregation with mutant HSCA2 (mtHSCA2) in combination with ISCU1 and/or SSR1. (*D*) Heat-induced CS aggregation with different amounts of SSR1. TAH1 is a TPR-containing yeast protein and used as a negative control that does not inhibit heat-induced CS aggregation. These test have been repeated 3 times.

### *ssr1-2* mutation causes mitochondrial dysfunction

HSCA2 and ISU1 are core components of the ISC pathway in Fe-S clusters biosynthesis and they have been well documented to interact with each other (Balk and Schaedler, 2014; Leaden et al., 2014). Additionally, our recent study on another weak SSR1 mutant allele *ssr1-1* indicated that SSR1 is important in regulating the function of mitochondrial electron-transport chain complexes, a few of which are well-known Fe-S cluster containing proteins (Han et al., 2021). We therefore hypothesized that the mitochondrial Fe-S cluster biosynthesis or the biosynthesis of Fe-S cluster containing proteins are impaired in our *ssr1-2* mutant.

We then investigated the activity and/or protein expression levels of some Fe-S proteins, namely mitochondrial complex I (CI) and complex II (CII), aconitase (ACO) and succinodehydrogenase 2 (SDH2), in *ssr1-2* seedlings. Given that aconitase is located in both mitochondria and cytosol, the aconitase activity from both compartments was measured. It was shown that the enzymatic activities of aconitase, CI, and CII as well as the protein level of SDH2 in *ssr1-2* were all dramatically lower than that in the wild type and the two double mutants, *ssr1-2 sus1* and *ssr1-2 sus2* (Fig. 7, A-C and E). As a control, the enzymatic activity of malate dehydrogenase, not an iron-sulfur protein, displayed no difference between the wild type and *ssr1-2* (Fig. 7D). Additionally, the protein level of ATP5A, a Fe-S cluster-independent subunit of complex V, was higher in *ssr1-2*, and ISU1 was comparable between the wild type, *ssr1-2*, and the double mutants (Fig. 7, D and E). Taken all data together, it is evident that SSR1 is required for Fe-S cluster containing proteins activity, thus further confirming the role of SSR1 in the maintenance of mitochondrial electron-transport chain as revealed by analyzing a weak mutant allele *ssr1-1* under the proline treatment (Han et al., 2021).

**Figure 7.**
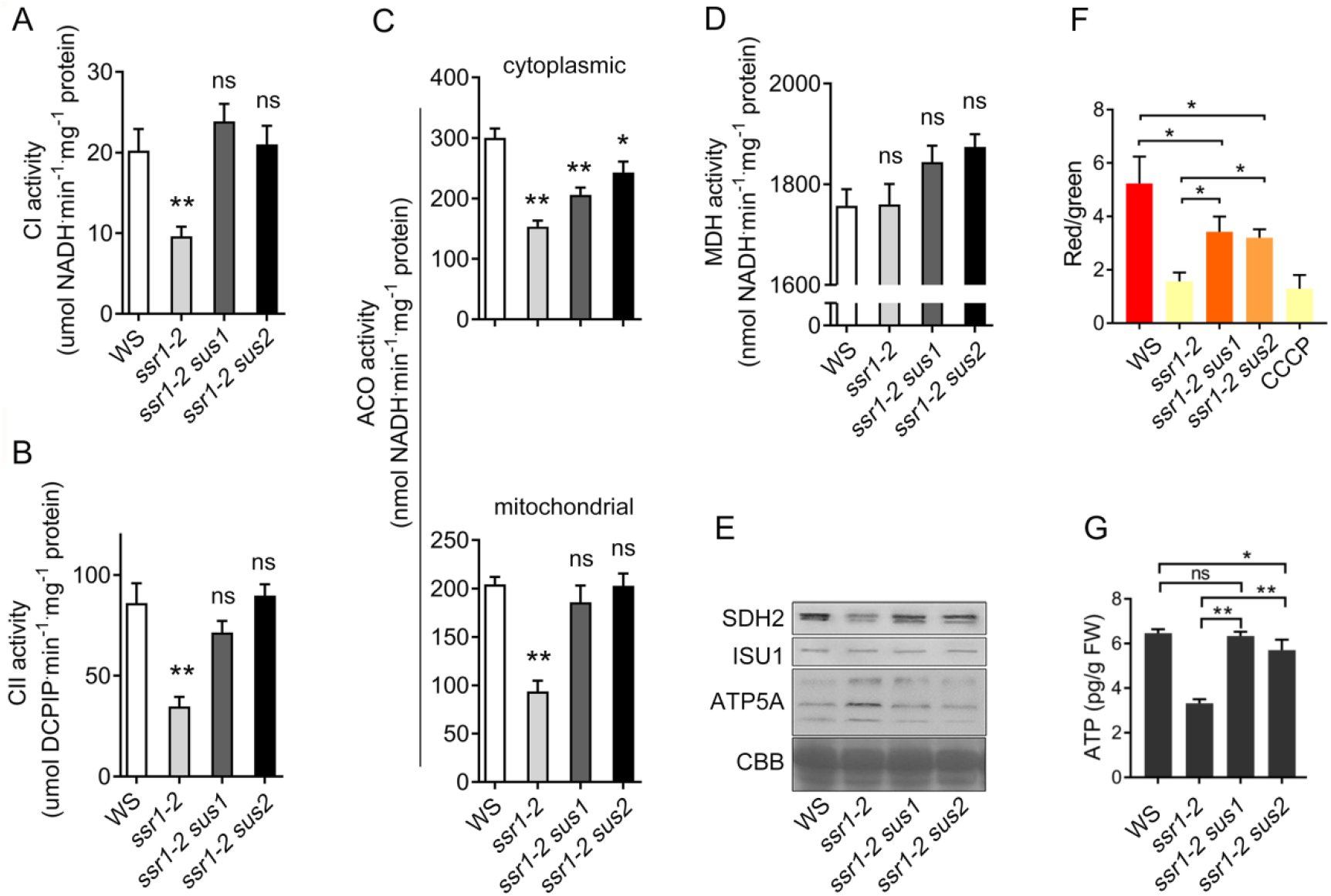
The activity and expression level of representative Fe-S containing enzymes, mitochondrial membrane potential and ATP contents decreased in *ssr1-2* mutant, and were restored with *sus1* or *sus2 suppressor*. Mitochondrial complex I (*A*), complex II (*B*), and aconitase (bottom panel of *C*) enzyme activities. Cytosolic aconitase (top panel of *C*) and malic dehydrogenase (*D*) enzyme activities. (*E*) Protein expression levels of SDH2, ISU, ATP5A as detected by corresponding antibodies and the total protein shown by Coomassie brilliant blue (CBB) staining. (*F*) Mitochondrial membrane potential (MMP). (*G*) Total cellular ATP contents. In (*F*), CCCP, a mitochondrial oxidative phosphorylation uncoupler, was used as positive control for mitochondria membrane depolarization. Flow cytometry was used for MMP detection and more than ten thousand of mitochondria in each sample were measured. Error bars represent standard deviation (*n*=3). ANOVA was used for statistical analysis (*F*=8.376 (*A*), *F*=13.76 (*B*), *F*=19.3 (top panel of *C*), *F*=17.34 (bottom panel of *C*), *F*=3.203 (*D*), *F*=80.33 (*G*), *DFn*=3; in (*F*), *F*=20.79, *DFn*=4). Asterisks indicate a significant difference from the WS or between two designated groups. \*\**P* < 0.01; \**P* < 0.05. “ns” represents statistically not significant.

Mitochondrial dysfunction often affects MMP and ATP content of cell. Thus, ATP content and mitochondrial membrane potential (MMP) were measured. To minimize the interference of chloroplasts, seedlings grown on MS medium in the dark were used for crude mitochondria preparation. Flow cytometry analysis showed that the MMP was greatly reduced in *ssr1-2* compared to the wild type, dropping to merely 30% of the wild type level, and was partially restored in *ssr1-2 sus1* and *ssr1-2 sus2* double mutants to approximate 55% of the wild type level (Fig. 7F). The content of ATP in *ssr1-2* was dramatically lower than that in the wild type and completely restored to the wild type level in *ssr1-2 sus1* and *ssr1-2 sus2* (Fig. 7G).

## Discussion

### SSR1 plays an important role in mitochondrial Fe-S cluster biosynthesis

SSR1 is a mitochondrial TPR domain-containing protein and was previously reported to be required for the function of mitochondria electron transport chain (Han et al., 2021) and primary root elongation (Zhang et al., 2015). However, the molecular mechanism underlying the function of *SSR1* remained elusive. In this report, we have pinpointed, by suppressor characterization, that the function of SSR1 in mitochondria as a crucial chaperone-like component to promote the association of HSCA2 with ISU1, facilitating the transfer of Fe/S cluster in ISC pathway.

It has been well documented in *E. coli* and yeast that the binding of HscA/Ssq1 to IscU/Isu1 is required for transferring Fe-S cluster from scaffold IscU/Isu1 to apo-protein (Hoff et al., 2003; Cupp-Vickery et al., 2004; Dutkiewicz et al., 2004; Kim et al., 2012). This regulatory system is evolutionarily conserved throughout higher organisms and has been recently identified in plant as well (Leaden et al., 2014; Armas et al., 2019; Armas et al., 2020). Here, our results illustrated that the optimal binding of HSCA2 to ISU1 is largely SSR1-dependent, since in the absence of SSR1 the binding between these two proteins is much weaker, as evidenced by in vitro pull-down assay (Fig. 4, C and D). However, HSCA2^G87D^ and ISU1^T55M^ could interact well with ISU1 and HSCA2, respectively, thus by-passing the need of SSR1. It appeared that the enhanced affinity between HSCA2 and ISU1 in two suppressor mutants is responsible for suppressing the short-root phenotype of *ssr1-2*, since artificially reducing the affinity between HSCA2 and ISU1^T55M^ by certain amino acid substitution in LPPVK motif of ISU1 would render ISU1^T55M^ incapable of suppressing *ssr1-2* phenotype (Fig. 5). Based on previous reports and our results, we proposed a model illustrating how SSR1 promotes the interaction between HSCA2 and ISU1 that is mediated by the HSCA2 LPPVK motif (Fig. 8).

**Figure 8.**
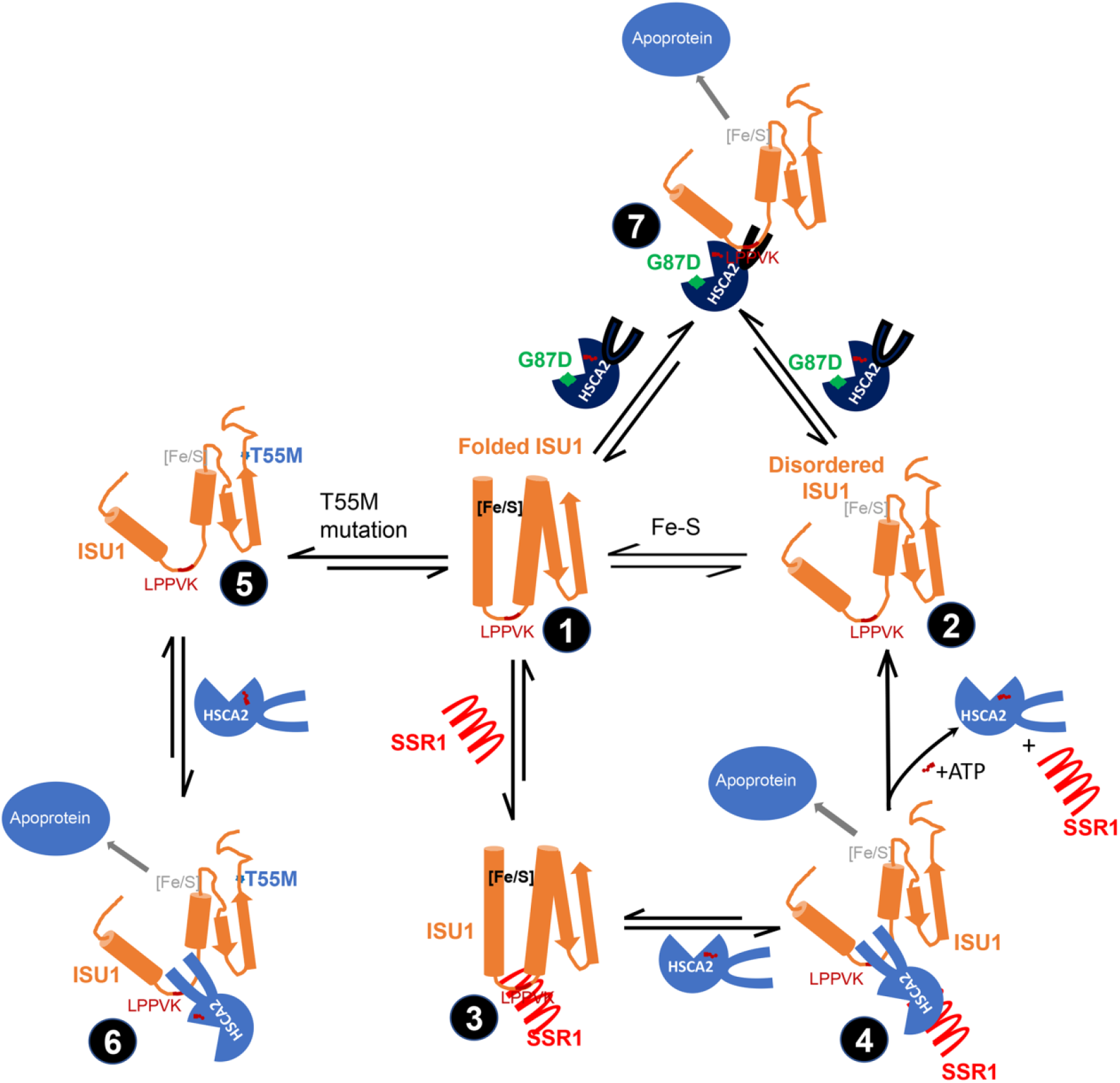
proposed model for HSCA2, ISCU1 and SSR1 interactions. Based on our results on suppressor characterization and previous reports on bacterial HSCA and ISU structural analysis, we propose that the free Arabidopsis ISU1 adopts two conformations in vivo, the disordered (state 1) and folded (state 2) structures. Fe-S cluster is assembled on disordered ISU1, and with loaded [4Fe-4S], ISU1 becomes folded. SSR1 binds folded ISU1 (state 3), and transfers ISU1 to HSCA2. HSCA2 binds and hydrolyzes ATP to ADP. ADP binding state of HSCA2 associates tightly with LPPVK and facilitates the formation of disordered ISU1 and [4Fe-4S] cluster transfer to apoproteins (state 4). Unlike ISU1, ISU1^T55M^ mutation (state 5) has higher affinity to HSCA2 in the absence of SSR1. Thus, ISU1^T55M^ has ability to transfer [4Fe-4S] to apoproteins, bypassing the help of SSR1 (state 6). Similarly, HSCA2^G87D^ mutation increases the binding affinity to ISU1 and bypasses the help of SSR1 in [4Fe-4S] transfer (state 7).

### SSR1 may function similarly as HscB/Jac1 in microorganisms

Iron-sulfur proteins play essential roles in many fundamental biological processes. Currently, it was estimated that about 100 iron-sulfur proteins are present in *Arabidopsis* (Balk and Pilon, 2011). Therefore, it was not surprising that *ssr1-2* mutant displayed pleiotropic growth retardation, with most striking phenotype being the short and swollen primary roots at seedling stage (Zhang et al., 2015). We have observed that *SSR1* knock-out mutation was correlated with altered expression patterns of a battery of auxin transport or response-related proteins (Zhang et al., 2015). This may well be associated with a shortage of Fe-S clusters supply, resulting in reduced activity of the aldehyde oxidase catalyzing the last step of the biosynthesis of auxin (Dai et al., 2005), as well as abscisic acid (Seo et al., 2000). Various degrees of growth retardation have been observed in other mutants deficient in the key components of Fe-S cluster assembly system. For example, ATM3, an ATP-binding cassette transporter in mitochondria, was proposed to transport some unknown intermediates for Fe-S cluster assembly in cytosol. *ATM3* T-DNA insertional mutant, *atm3-1*, showed dramatically reduced content of abscisic acid and the enzymatic activities of some cytosolic Fe-S proteins, resulting in a pleiotropic growth defect, such as small stature, lower chloroplast numbers and male sterility (Bernard et al., 2009; Teschner et al., 2010). Smaller plantlet size and bushy phenotype were also observed in Arabidopsis lines with knocking-down of *AtNFS1* and *AtISU1* (Frazzon et al., 2007).

In a T-DNA insertional mutant of *AtHSCB* with undetectable full-length transcript and protein, however, no above-mentioned growth defect was observed, despite greatly reduced activities of Fe-S enzymes aconitase and succinate dehydrogenase (Xu et al., 2009; Leaden et al., 2016). It was well-established in microorganisms that HscB/Jac1, a homolog of AtHSCB, could promote the interaction between HscA/Ssq1 and IscU/Isu1 and stimulate the ATPase activity of HscA/Ssq1, which is required for the transfer of Fe-S clusters (Silberg et al., 2004; Chandramouli and Johnson, 2006; Francesco et al., 2008). The *jac1* yeast mutant is lethal, indicating that *JAC1* is an essential gene (Voisine et al., 2001). The relative mild phenotype of *athscb* mutant (Xu et al., 2009; Leaden et al., 2016) would suggest a possible existence of functional redundancy for *AtHSCB*. Our results demonstrated that SSR1 could interact with both HSCA2 and ISU1, and the presence of SSR1 is required for the effective interaction between HSCA2 and ISU1 (Fig. 4 and Fig. 5), suggesting that SSR1 could partly mimic the function of HscB/Jac1 in Fe-S cluster biosynthesis in mitochondria, though their mechanism of action may be different. In addition, it seems that SSR1 plays much more important role in ISC pathway than does HscB, since the growth defect of *SSR1* loss-of-function mutant is so dramatic.

It should be noted that in bacteria HscB did not exhibit intrinsic chaperone activity and only function as a cochaperone for HscA (Silberg et al., 1998). However, we have shown here that SSR1, which does not have typical J-domain, displayed strong chaperone activity that was far more powerful than that of HSCA2. Whether this may be responsible for the difference between SSR1 and HscB in their function in ISC pathway await further investigation.

### SSR1 may have evolved as a stress-adapting component for ISC system in Plant kingdom

Unlike most of the other components in ISC system, *SSR1* is not inherited from microorganism. Rather, it emerged as a new gene present only in plant kingdom with a distant homolog appearing already in green algae (Supplemental Fig. S2). Currently, we are not clear what the evolutionary advantage for plant to possess *SSR1* is. One clue comes from our characterization of *ssr1-1*, a weak allele with a truncated SSR1 protein. It has been shown that *ssr1-1* is hypersensitive to osmotic stress (Zhang et al., 2015) and proline treatment (Han et al., 2021), both of which were believed to cause electron overflow in mitochondrial electron transport chain (mETC) due to inhibited mETC activity or elevated electron supply (Phang et al., 2008; Launay et al., 2019). Actually, ISC machinery is prone to inhibition by ROS under various environmental stresses (Liang et al., 2014) during their sessile life cycle, to which plants must respond and adapt. Therefore, we propose that *SSR1* may be required for protecting ISC machinery from stresses via stabilizing the interaction of HSCA2 and ISU1, which is crucial for supplying Fe-S cofactor to many components in mETC. In supporting our proposition, *SSR1* was shown to be transcriptionally up-regulated by several abiotic stresses (Supplemental Fig. S4).

In conclusion, we have presented evidences that SSR1 is a new and vital chaperone-like component of ISC pathway specifically for Fe-S clusters biosynthesis in plants.

## Materials and methods

### Plant materials and growth conditions

For seedling growth experiments, surface-sterilized seeds of *Arabidopsis* wild type (WS), transgenic plants and mutants were sown on solid Murashige and Skoog (MS) medium, containing 1% sucrose and 0.25% phytogel, and stratified at 4°C for 2 days in the dark before being transferred to an incubator for germination at 22-24°C,16h light/8h dark photoperiod with light an intensity of 110 μmole.m^-2^.sec^-1^. To grow to maturity, 4-day-old seedlings were planted in soil and grown under similar condition as in the incubator. All transgenic plants used in this study were listed in Table S2. Primers used for real-time qPCR and T-DNA insertion mutant detection were listed in Table S3.

### Cloning of *SUS1* and *SUS2*

*ssr1-2* mutants harboring *sus1* or *sus2* suppressor that were originally generated by ethyl methanesulfonate mutagenesis were backcrossed twice with *ssr1-2* parental line to obtain F1 plants. The F1 plant was self-pollinated to obtain F2 progeny, which were further planted and harvested individually. From the F3 population, two pools were formed, with each containing 30 homozygous lines (no segregation for the root-length phenotype) with either the longest roots or the shortest roots. DNAs were isolated from *ssr1-2* and two pools and were subjected to genome re-sequencing. Super Bulked Segregant Analysis (Turner et al., 2010) was used to screen for candidate mutation sites that are correlated with the root length phenotype.

### Plasmids Construction and plant transformation

All constructs and primers used in this study were listed in Table S4. Briefly, for genetic complementation, the genomic sequences for *sus1, sus2, sus5* and *sus6*, which encode HSCA2^G87D^, ISU1^T55M^, ISU1^G106D^ and ISU1^A143T^ respectively, were amplified and cloned from corresponding suppressor mutants. To generate HSCA1^G82D^ from wild type HSCA1, the *HSCA1* coding sequence (CDS) was amplified in two halves separately, and then fused together by overlap extension PCR with the simultaneous introduction of the mutation. All complementation constructs were based on the binary vector pCAMBIA1300 and pCAMBIA1300-super-Flag. For tissue-specific expression analysis with GUS reporter gene, the promoter of *HSCA2* was amplified and inserted into pCAMBIA1391. For subcellular localization analysis, the CDS of *HSCA2* was amplified and cloned into pBI121-mCherry. The reporter constructs pSSR1-GUS and pSSR1-GFP have been described previously (Zhang et al., 2015). For protein expression in *E. coli*, the coding sequences of *HSCA2, SSR1* and *ISU1* were cloned into vector pRSET with cMyc Tag. Amino acid substitution mutants around the LPPVK motif of ISU1 were also generated with overlap extension PCR. To generate constructs, transfected and transgenic plants for co-IP analysis, the promoter and CDS of *HSCA2* was amplified and first inserted into pBI121-cMyc. Then the Myc-tagged expression cassette were amplified and inserted into pCAMBIA1300. The pSSR1-Flag was described previously (Zhang et al., 2015). For BiFC assays, the CDS of *SSR1, HSCA2* and *ISU1* from wild type were inserted into pE3242 or pE3228 to generate either nVenus tagged or cCFP-tagged constructs, respectively. See Lee’s work for more details about vectors pE3228 and pE3242 (Lee et al., 2008).

Whenever needed, *Arabidopsis* plants were transformed with Agrobacteria-mediated floral dipping method(Clough and Bent, 1998) or the protoplasts were transfected with expression constructs as described (Yoo et al., 2007). The trangenic plants were screened on kanamycin- or hygromycin B-containing MS medium depending on the vector used. The integration of the transgene was confirmed by PCR.

### Protein expression and purification from *E. coli*

His_6_-tagged protein expression and purification from *E. coil* was carried out as described previously (Leaden et al., 2014). Briefly, BL21(DE3) bacterial strains with respective constructs were cultured in LB liquid medium at 37°C to OD_600nm_≈0.5, and then induced with IPTG at a final concentration of 1mM for 6 h at 28°C. Cells were harvested, re-suspended in buffer A [20 mM Tris–HCl, 200 mM NaCl, 30 mM imidazole and 1 mM phenylmethylsulfonyl fluoride (PMSF), pH 7.4] and then disrupted by sonication. The suspensions were centrifuged at 10,000 ×g for 15 min at 4°C. The supernatants of the His_6_-tagged proteins obtained were incubated with 500μL Ni Sepharose (GE Healthcare 17-5318-06) and then washed twice with buffer A. The recombinant proteins were eluted with buffer B (500 mM imidazole in buffer A). For proteins to be used for in vitro chaperone activity assays, the eluants were further applied to size exclusion chromatography with the Superdex 75 or Superdex 200 column with ÄKTA Purifier 10 FPLC system (GE Healthcare). The *E. coli* extracts containing the recombinant proteins without His_6_-tag were used for the pull-down experiments.

### Protein-protein interaction assays

Pull-down assay was carried out as described previously (Leaden et al., 2014). Briefly, about 10 ug purified His_6_-tagged HSCA2, HSCA2^G87D^ or SSR1 was incubated with BL21(DE3) cell extracts expressing SSR1, ISU1 or other mutant ISU1 without His_6_-tag for 1 hour at 4°C. Next, 20 μL Ni Sepharose pre-equilibrated with buffer A as described in previous section, were added and incubated for another 2 hours at 4°C. After washing twice with buffer A, the proteins binding to Sepharose were eluted by appropriate volume of buffer B, and analyzed by Western blot.

BiFC assays were performed via transient expression in Arabidopsis mesophyll protoplasts as previously described(Yoo et al., 2007). Co-IP assays from transfected protoplasts were performed with C-Myc Isolation Kit (130-091-123) and DYKDDDDK (Flag) Isolation Kit (130-101-591) from Miltenyi Biotec. Briefly, 12-day-old transgenic seedlings containing *HSCA2*_*pro*_*:HSCA2-Myc* and *35S*_*pro*_*:SSR1-Flag* were ground to fine powder in liquid nitrogen. About 500 mg powder was transferred to 2 mL Eppendorf tube, before 1.5 mL pre-cooled lysis buffer (50 mM Tris-HCl, pH7.9, 120 mM NaCl, 10% glycerol, 10 mM DTT, 5 mM EDTA, 1% PVP, 1% NP-40, 1 mM PMSF, protease inhibitor cocktail) was added and mixed well with the powder. The homogenates were incubated for 30 minutes on ice with occasional mixing and then centrifuged for 10 minutes at 10,000 ×g at 4°C to get rid of the debris. Total input samples were taken from the supernatant, and the remaining supernatant was transferred to a fresh 1.5 mL tube with 50 μL antibody and was incubated on ice for 40 minutes. Place μ column (130-042-701) in the magnetic field of the μMACS Separator and prepare the μ column by applying 200 μL lysis buffer on the column. Transfer the supernatant into the μ column and let the lysate run through. Rinse the column with 4×200 μL of lysis buffer and 1×100 μL wash buffer (20 mM Tris-HCl, pH 7.5). Pre-heated elution buffer (from Kit) was subsequently added to elute the immunoprecipitated, which was used for Western blot.

The antibodies used for Western blot were purchased from Abcam (anti-ISU: ab154060; anti-SDH2: ab154974; anti-ATP5A: ab14748), Tiangen Biotech (anti-His: AB102) and Sigma-Aldrich (anti-Myc: M4439; anti-Flag: F3165).

### In vitro chaperone activity assay

All tested proteins and citrate synthase (CS, Sigma, C3260) was dialyzed in 20 mM HEPES-KOH, pH 7.5, 150 mM KCl, 10 mM MgCl_2_ before being used for the heat-induced aggregation assay. CS (500 nM) was prepared in a final volume of 150 μL 20 mM HEPES-KOH (pH 7.5), 2.8 mM β-mecaptoethanol with different amounts of tested proteins. The mixtures were added to a 96-well microplate and heated at 45°C. Light scattering at 340 nm was monitored at 45°C in a Synergy 4 spectrophotometer (BioTek) for 90min. Control measurements were performed with purified test proteins alone in the absence of CS and a His_6_-tagged yeast TPR-containing Tah1 protein expressed and purified from *E. coli* was also used as a negative control (Zhao et al., 2008).

### Isolation of mitochondria and mitochondrial functionality assays

Mitochondria isolation was performed essentially as previously described (Han et al., 2021) from 10-day-old seedlings grown in MS medium and in the dark. Crude mitochondria were used for mitochondrial membrane potential (MMP) analysis, and respective enzymatic activity assays focusing on complex I (CI), complex II (CII), aconitase (ACO), and malate dehydrogenase (MDH). The enzymatic activities of CI, CII, ACO and MDH were detected with kits from Suzhou Comin Biotechnology Co. Ltd (FHTA-2-Y, FHTB-2-Y, ACO-2-Z, NMDH-2-Y). Cellular ATP contents were also measured for seedlings grown in the dark as MMP analysis. ATP content and MMP detection were analyzed by kits from Beyotime (S0026, C2006). JC-1, a fluorescent indicator, was used to measure the MMP. Flow cytometry (Beckman: Moflo XDP) was used to detect the fluorescence in two channels (red and green) and more than ten thousand of mitochondria in each sample were measured.

### Searching homologous proteins of SSR1 in other species

The amino acid sequence of *Arabidopsis* SSR1 was used to search for homologous proteins from microorganism to higher organisms using UniPro BLAST and NCBI DELTA-BLAST. The similarity of ISC components between *E. coli* and *Arabidopsis* (ISU1/2/3, HSCA1/2, HSCB, ADX1/2 and FH) was used as a reference for threshold value (*E* value less than 1*10^−4^; identity larger than 20%) to filter out the returned proteins displaying low similarity to SSR1. Then, the candidates from different representative species were used as query to search in *Arabidopsis* database with the same threshold value. If *Arabidopsis* SSR1 was returned as subject, the candidate would be selected as homologous protein.

Homologous proteins from representative species at different evolutionary status were used to construct a phylogenetic tree by using MEGA7 (Kumar et al., 2016). The evolutionary history was inferred using the Neighbor-Joining method. The percentage of replicate trees in which the associated taxa clustered together in the bootstrap test (1,000 replicates) is shown next to the branches.

### Accession Numbers

Sequence data for genes and proteins presented in this article can be found in the Arabidopsis Genome Initiative of GenBank/EMBL database under the following accession numbers: *SSR1* (AT5G02130), *HSCA2* or *mtHSC70-2* (At5g09590), *ISU1* (At4g22220), *HSCA1* or *mtHSC70-1* (AT4G37910), *Actin 7* (AT5G09810).

## Acknowledgements

We thank Prof. Dr. Hongzhi Kong from Institute of Botany, Chinese Academy of Sciences for the valuable suggestions on phylogenetic tree construction, and Bona Mu for her help in the large-scale protein purification from *E. coli*.

---

**Supplemental Figure S1.**
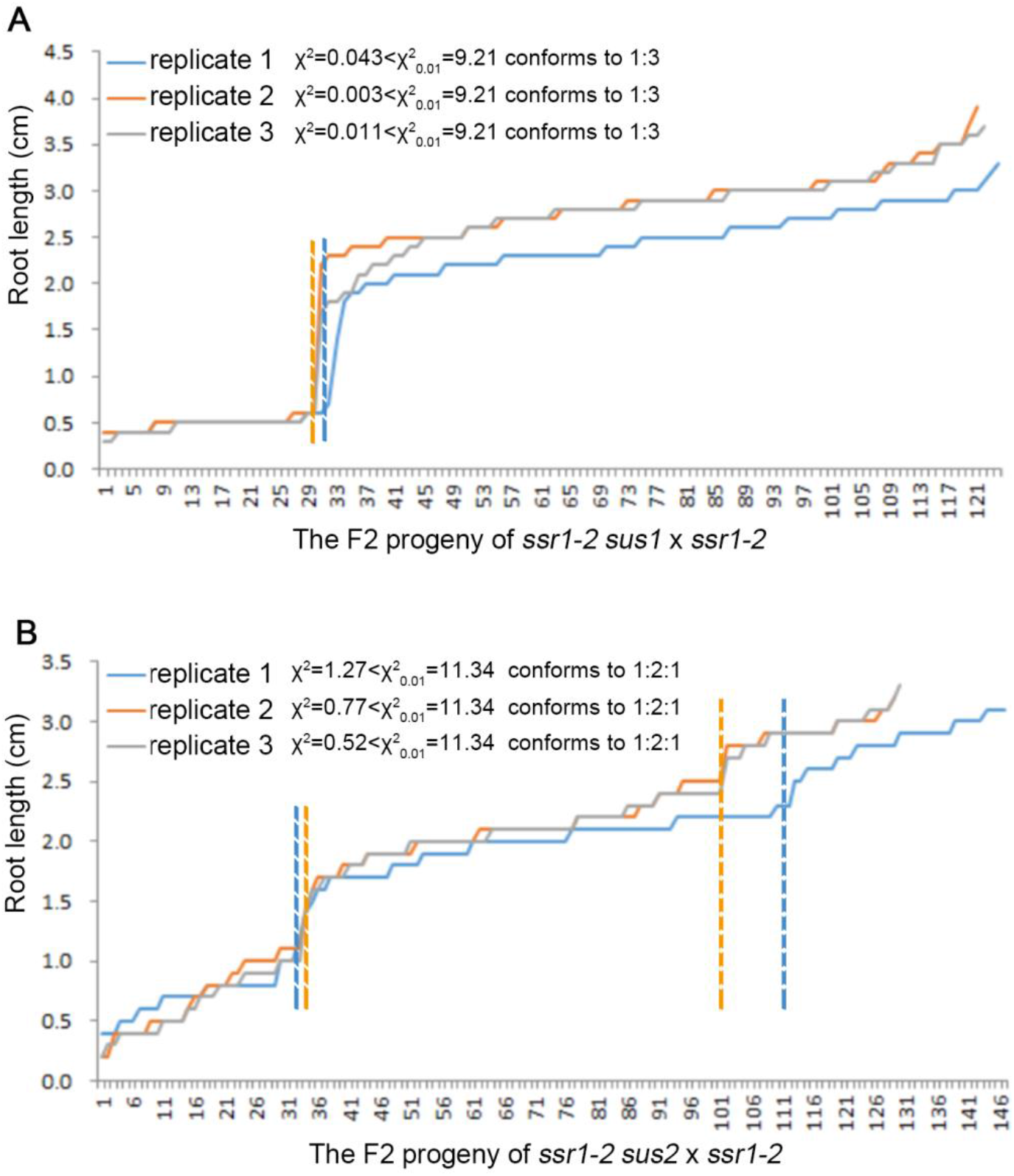
Root length of the F2 population seedlings from backcrossed suppressor mutant lines. The two suppressor lines *ssr1-2 sus1* (*A*) and *ssr1-2 sus2* (*B*) were backcrossed with *ssr1-2* and resulting F1 plants were self-pollinated to generate F2 seeds. The primary root length of the F2 seedlings grown at 10-days-old were measured and graphed. The original root length data are shown in Table S1. Three biological repeats were performed and shown in differently colored lines.

**Supplemental Figure S2.**
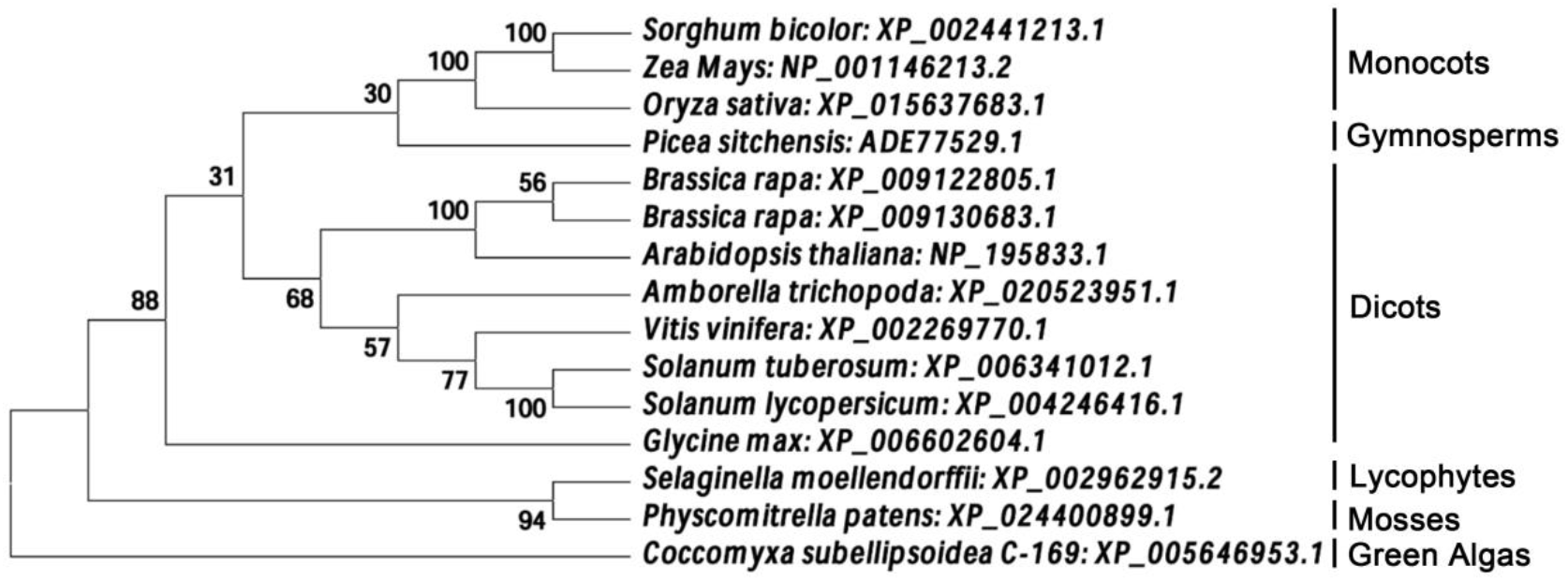
SSR1 exists only in plant kingdom. Homologous proteins of SSR1 were identified in green alga, basal and higher plants, but not in microorganism and animalia. Homologous proteins from representative species at different evolutionary status were used to construct a phylogenetic tree by using MEGA7. The evolutionary history was inferred using the Neighbor-Joining method. The percentage of replicate trees in which the associated taxa clustered together in the bootstrap test (1000 replicates) are shown next to the branches. See method for more details.

**Supplemental Figure S3.**
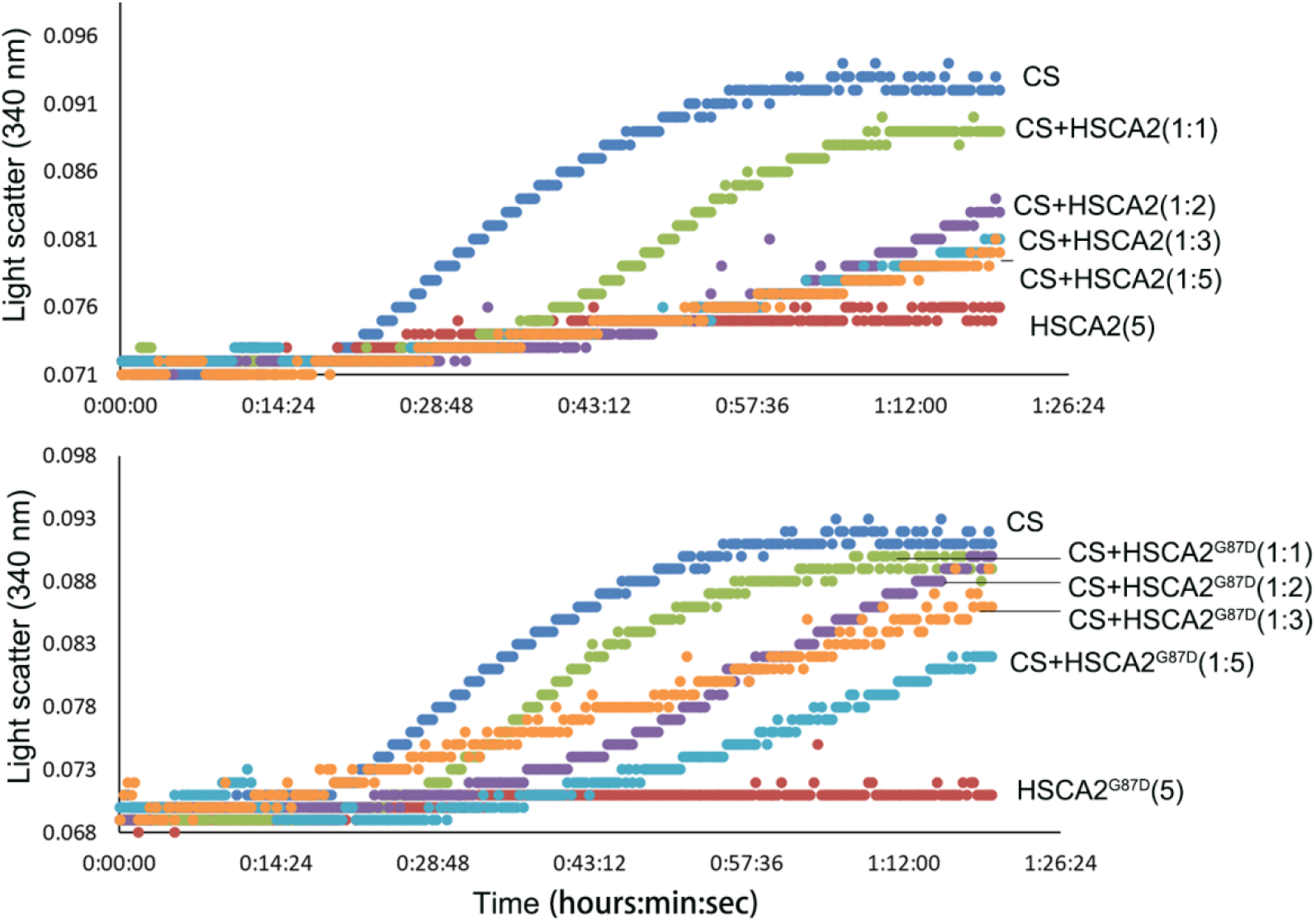
General chaperone activity assays of purified HSCA2 and HSCA2^G87D^. Heat-induced aggregation of citrate synthase (CS) was performed at 45°C for 90min with different amount of purified test proteins. The molecular ratios of CS to tested proteins are indicated following each protein sample.

**Supplemental Figure S4.**
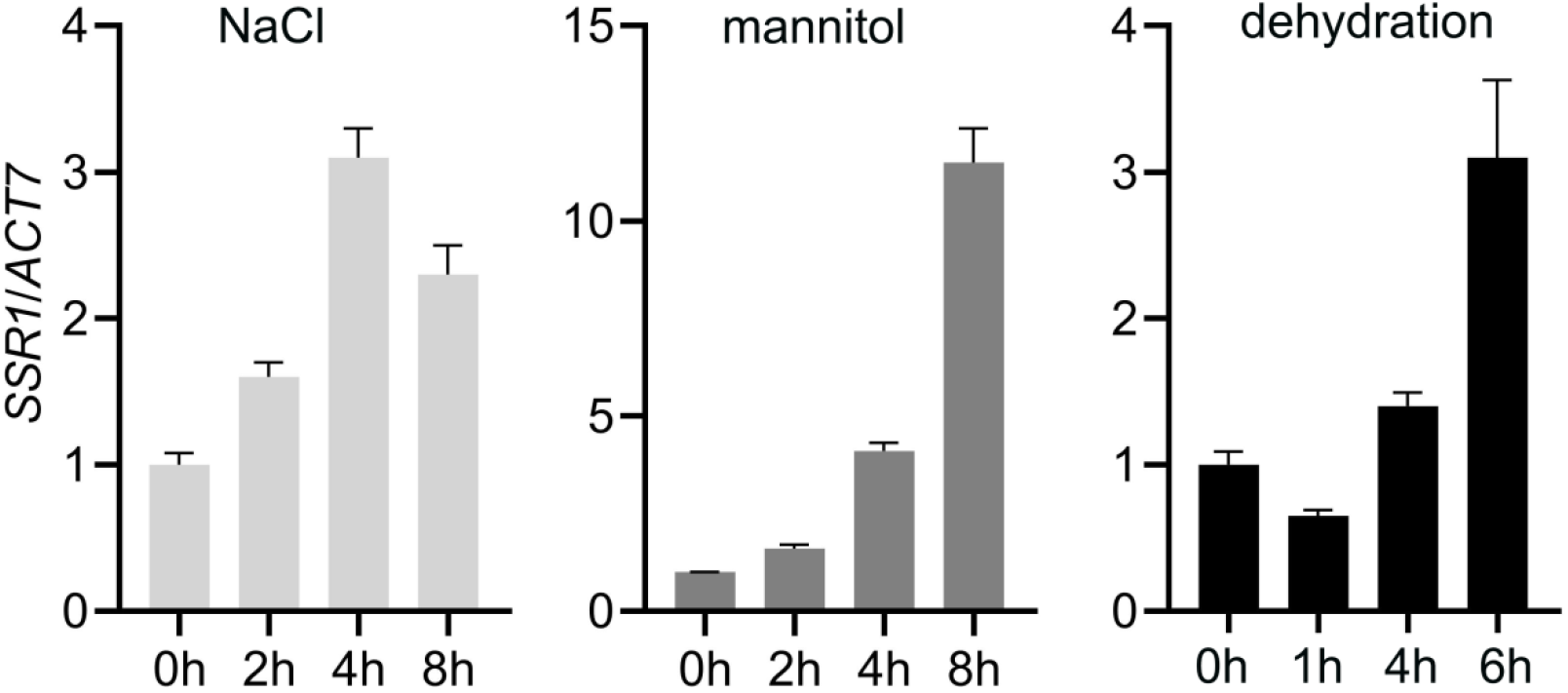
Relative expression of *SSR1* upon different stresses. *Actin 7* was used as internal control. Wild type WS seedlings grown at 10-days-old were treated with 100 mM NaCl, 150 mM Mannitol and 20-day-old seedlings were removed from soil for dehydration treatment. Samples were taken at different time after treatment for total RNA extraction. The primers for *ACT7* genes were used as an internal control. Three biological repeats were performed and the representative result was shown.

**Supplemental Table S1**. Root length of backcrossed F2 seedlings.

**Supplemental Table S2**. Tansgenic plants used in this study.

**Supplemental Table S3**. Primers for real-time qPCR or plant lines genotyping.

**Supplemental Table S4**. Plasmid constructs and primers used in this study.

